# Earth-Gravity Congruent Motion Facilitates Ocular Control for Pursuit of Parabolic Trajectories

**DOI:** 10.1101/547497

**Authors:** Björn Jörges, Joan López-Moliner

## Abstract

There is evidence that humans rely on an earth gravity (9.81 m/s^2^) prior for a series of tasks involving perception and action, the reason being that gravity helps predict future positions of moving objects. Eye-movements in turn are partially guided by predictions about observed motion. Thus, the question arises whether knowledge about gravity is also used to guide eye-movements: If humans rely on a representation of earth gravity for the control of eye movements, earth-gravity-congruent motion should elicit improved visual pursuit. In a pre-registered experiment, we presented participants (n=10) with parabolic motion governed by six different gravities (−1/0.7/0.85/1/1.15/1.3g), two initial vertical velocities and two initial horizontal velocities in a 3D environment. Participants were instructed to follow the target with their eyes. We tracked their gaze and computed the visual gain (velocity of the eyes divided by velocity of the target) as proxy for the quality of pursuit. An LMM analysis with gravity condition as fixed effect and intercepts varying per subject showed that the gain was lower for −1g than for 1g (by-0.13, SE=0.005). This model was significantly better than a null model without gravity as fixed effect (p<0.001), supporting our hypothesis. A comparison of 1g and the remaining gravity conditions revealed that 1.15g (by 0.043, SE=0.005) and 1.3g (by 0.065, SE=0.005) were associated with lower gains, while 0.7g (by 0.054, SE=0.005) and 0.85g (by 0.029, SE=0.005) were associated with higher gains. This model was again significantly better than a null model (p<0.001), contradicting our hypothesis. Post-hoc analyses reveal that confounds in the 0.7/0.85/1/1.15/1.3g condition may be responsible for these contradicting results. Despite these discrepancies, our data thus provide some support for the hypothesis that internalized knowledge about earth gravity guides eye movements.

## Introduction

There is a growing corpus of research indicating that humans use their knowledge of earth gravity in a broad range of tasks, such as grasping^1^, catching and interception^2–11^, duration estimation^12^, the perception of biological motion^13–15^ and visual estimation of motion^16^, while arbitrary accelerations are generally neglected^17,18^ or used insufficiently. Humans seem to rely on their knowledge of earth gravity to predict target motion in space and/or time. This predictive information is then combined with online information in order to estimate relevant variables of the environment (such as time-to-contact or location of interception). On the other hand, eye movements have been shown to be partially guided by predictive mechanisms^19,20^. If gaze behavior is influenced by predictions and an “internal model of gravity”^4^ or “earth gravity prior”^21^ drives prediction in many different computations, then this internal representation of 1g should allow humans to follow targets more closely when their trajectories are governed by earth gravity. And indeed, this line of reasoning has recently been brought into focus: Delle Monache et al^22^ presented participants with targets moving along parabolic trajectories that were generally governed by earth gravity. In the second half, a perturbation with 0g or 2g could occur. The trajectories were presented in a 2D scene with pictorial cues. Participants tracked targets less successfully during the 0g perturbations than during 2g perturbations or trials without perturbation, which can be taken as some evidence for the role of a representation of earth gravity being used in gaze behavior. While this should, in principle, also impact gaze following during the 2g perturbations, the 2D presentation may have affected the recruitment of an internal model of gravity negatively^12,23^. While ocular movements have been studied for parabolic 1g motion^24–28^, to our knowledge, human gaze behavior has not been explored for parabolic motion governed by uniformly non-terrestrial gravities. This is of special interest because smooth pursuit of curved trajectories is a demanding task that requires a complex interplay of ocular muscles. It would thus be particularly beneficial to employ all available information, including a strong prior for earth gravity, to plan and execute these movements.

Based on the above considerations, the present paper tests the hypothesis that participants should follow targets on parabolic trajectories more successfully when they are governed by earth gravity rather than by earth-discrepant gravities. Furthermore, we tested whether we can replicate previous results that knowledge of earth gravity is used to extrapolate motion information for occluded stimuli.

Both hypotheses have been pre-registered at the Open Science Foundation (https://osf.io/8vg95/), and any other, exploratory analyses of our data are clearly marked as such and should be treated with the due caution.

## Methods

### Participants

A total of eleven (n = 10) participants performed the task, among which one of the authors (BJ). All had normal or corrected-to-normal vision. The remaining participants were in an age range of 23 and 34 years and five (n = 3) were female. We did not test their explicit knowledge of physics, as previous studies suggest that explicit knowledge about gravity has no effect on performance in related tasks^29,30^. All participants gave their informed consent. The research in this study is part of an ongoing research program that has been approved by the local ethics committee of the University of Barcelona. The experiment was conducted in accordance with the Code of Ethics of the World Medical Association (Declaration of Helsinki).

### Stimuli

We presented participants with targets of tennis ball size (radius = 0.033 m), shape and texture that moved on parabolic trajectories. The trajectories were determined by the gravity levels (0.7,0.85,1,1.15,1.3g,-1g), the initial vertical velocities (4.5 and 6 m/s) and the initial horizontal velocities (3 and 4 m/s). The different kinetic profiles, as well as the occlusion condition (Short Occlusion: last 20-25%; Long Occlusion: last 45-50% of the trajectory), were presented in random order, but the method guaranteed that each combination was presented the same amount of times. The parabolas were presented in the fronto-parallel plane with no change in depth. Air resistance was simulated to provide a more realistic stimulus. The following equations (http://www.demonstrations.wolfram.com/ProjectileWithAirDrag/) determine the x position of the target in time *x(t)*, and the y position of the target in time *y(t)*, respectively, including air resistance:

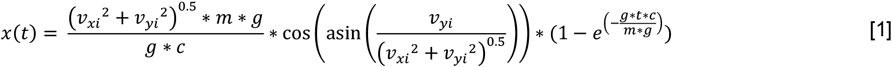

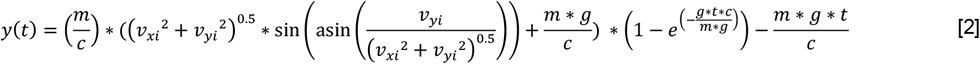

With *ν*_*xi*_ being the initial horizontal velocity, *ν*_*yi*_ the initial vertical velocity, *m* the mass of the target (0.057 kg), *g* the respective gravity value and *c* being the drag coefficient, where we chose 0.005.

For description of the parabolas, we use a coordinate system where the observer’s position is defined as x = y = z = 0; the x axis runs from left to right, the y axis from down to up and the z axis away from the observer in depth. The starting y position was half a meter above the ground (y = 0.5 m) for positive gravity values (0.7-1.3g) and 3.5 m for negative gravity values (−1g), while the starting x position was moved to the left from the middle of the scene by half of the overall length of the trajectory (x = -length/2 m). The target travels to the right, such that the peak of the parabola was always reached at x = 0 m (or the lowest point for the inverted parabolas). The ball’s z position remained constant at z = −6.15 m. The target disappeared at a random point between 20% and 25% (Short Occlusion) or 45% and 50% (Long Occlusion) of the time it would take for it to return to the initial height (y = 0.5 m or y = 3.5 m, respectively). The y end position was marked with an elongated table that was displayed in the target area of the room for targets with positive gravities; it was marked with an elongated lamp hanging from the ceiling for inverted stimuli. We presented the trajectories in a rich environment that provided 3D cues about the object’s position in depth (see Figure 1) and used a known object (a tennis ball) as target to recruit prior knowledge consistent with the geometry on display. This has been shown to help activate the internal model of gravity^10,23,26^, that we have previously suggested to be an earth gravity prior^21^. This environment was constructed such that no low-level cues such as differences in brightness and contrast with the target differed significantly between the different trajectories.

**Figure 1:**
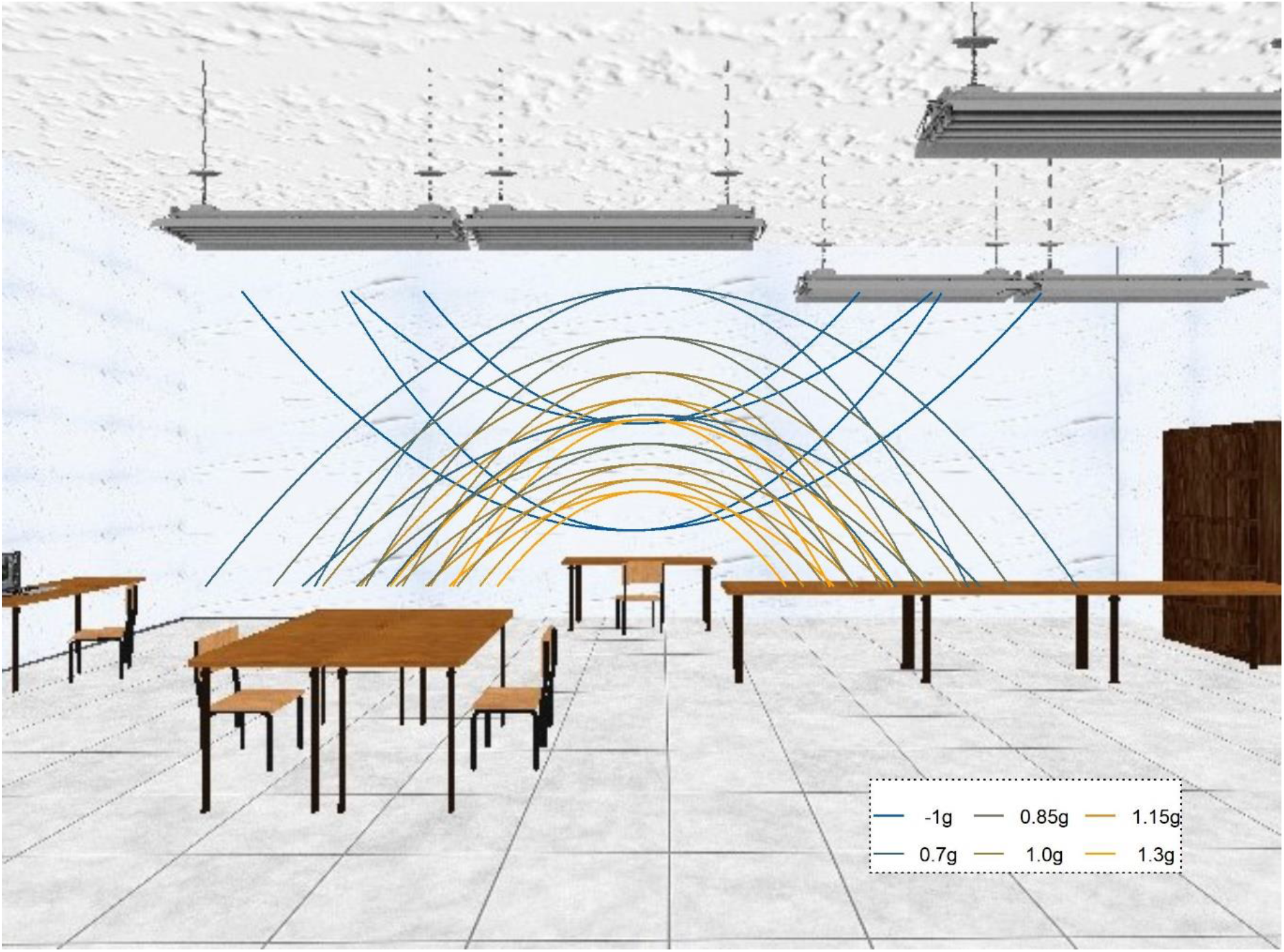
2D depiction of the visual scene used as environment for stimulus presentation. The stimulus was always presented in front of the white wall and never crossed other areas (such as the lamps of tables) that could introduce low level differences in contrast etc. The lines denote the different parabolic trajectories that along which the targets travelled.

### Apparatus

Two Sony laser projectors (VPL-FHZ57) were used to provide overlaid images on a back-projection screen (244 cm height and 184 cm width) with a resolution of 1920×1080 pixels. The frequency of refresh of the image was 85 Hz for each eye. Circular polarizing filters were used to provide stereoscopic images. Participants stood at 2 m distance centrally in front of the screen and used polarized glasses to achieve stereoscopic vision of the visual scene and the target. The shown disparity was adapted to each participant’s inter-ocular distance. Eye-movements were recorded with a Pupil Labs eye-tracking device working at 200 Hz^31^. We employed the surface-tracker technology provided in the Pupil Labs Software to map eye-tracking data directly to the real-world coordinates of our display. The pupil labs eye-tracker achieves generally an accuracy of below 1° visual degrees (see pupil labs documentation under https://docs.pupil-labs.com/#notes-on-calibration-accuracy). In our setup, we achieved accuracies of < 0.5° for most of our participants; whenever the calibration was > 1°, we adjusted the cameras and repeated the calibration. The stimuli were programmed in PsychoPy^32^; we added the code to our pre-registration (https://osf.io/8vg95/). The projectors introduced a delay of 0.049259 s (SD = 0.001894 s) that will be accounted for in the analysis of timing responses.

### Procedure

Participants were first instructed to pursue the target closely with their eyes and to indicate via mouse button click when they believed the target had returned to the starting level (y = 0.5 m/y = 3.5 m). We familiarized subjects with 48 training trials (each combination of experimental variables once), in which the ball reappeared upon mouse click, thus indicating the spatial error. Then, we calibrated the eye-tracker and started data collection. We presented the stimuli in four blocks: 3 blocks of 320 trials each (5 gravities from 0.7g to 1.3g; 2 initial vertical velocities; 2 initial horizontal velocities; 2 occlusion conditions; 8 repetitions per combination), and one block of 384 trials (2 gravities – 1g and −1g –; 2 initial vertical velocities; 2 initial horizontal velocities; 2 occlusion conditions; 24 repetitions per combination) for a total of 1344 trials. After each block, participants could take a break, after which the calibration of the eye-tracker was repeated to avoid that potential displacement of the eye-tracker on the subjects’ head contaminated the data. Five subjects (s1, s3, s5, s7, s9) received the 1g/-1g block as first block, while the other five subjects (s2, s4, s6, s8, s10) received it as last block.

### Justification of Sample Size

The relative contribution of participants and number of observations to overall power differs according to the ratio of inter-subject variability and intra-subject variability^33^. We assume that the effect is relatively homogeneous between subjects, with variability thus stemming mostly from trial-to-trial intra-subject differences. Figure 2 visualizes this relationship for two effect sizes (a very small 0.05 difference in gain versus a small 0.1 difference in gain), while assuming that intra-subject variability is 6 times higher than inter-subject variability. A combination of about 60 observations per condition and 10 subjects should thus provide sufficient power to capture any relevant effect. However, in order to power potential post-hoc analyses for possible interactions between gravities and initial speeds, we choose to repeat each combination of 2 vertical velocities, 2 horizontal velocities, 2 occlusion conditions and 5 gravities 24 times, making for a total of 2*2*2*5*24 = 960 trials. We compare 1g and −1g parabolas in a separate block, again with 2 vertical velocities, 2 horizontal velocities and 2 occlusion conditions, which adds another 2*2*2*2*24 = 384 trials. This way, we also satisfy the generic recommendation of total observations per condition issued by Brysbeard & Steven^34^ in order to achieve a properly powered experiment.

**Figure 2:**
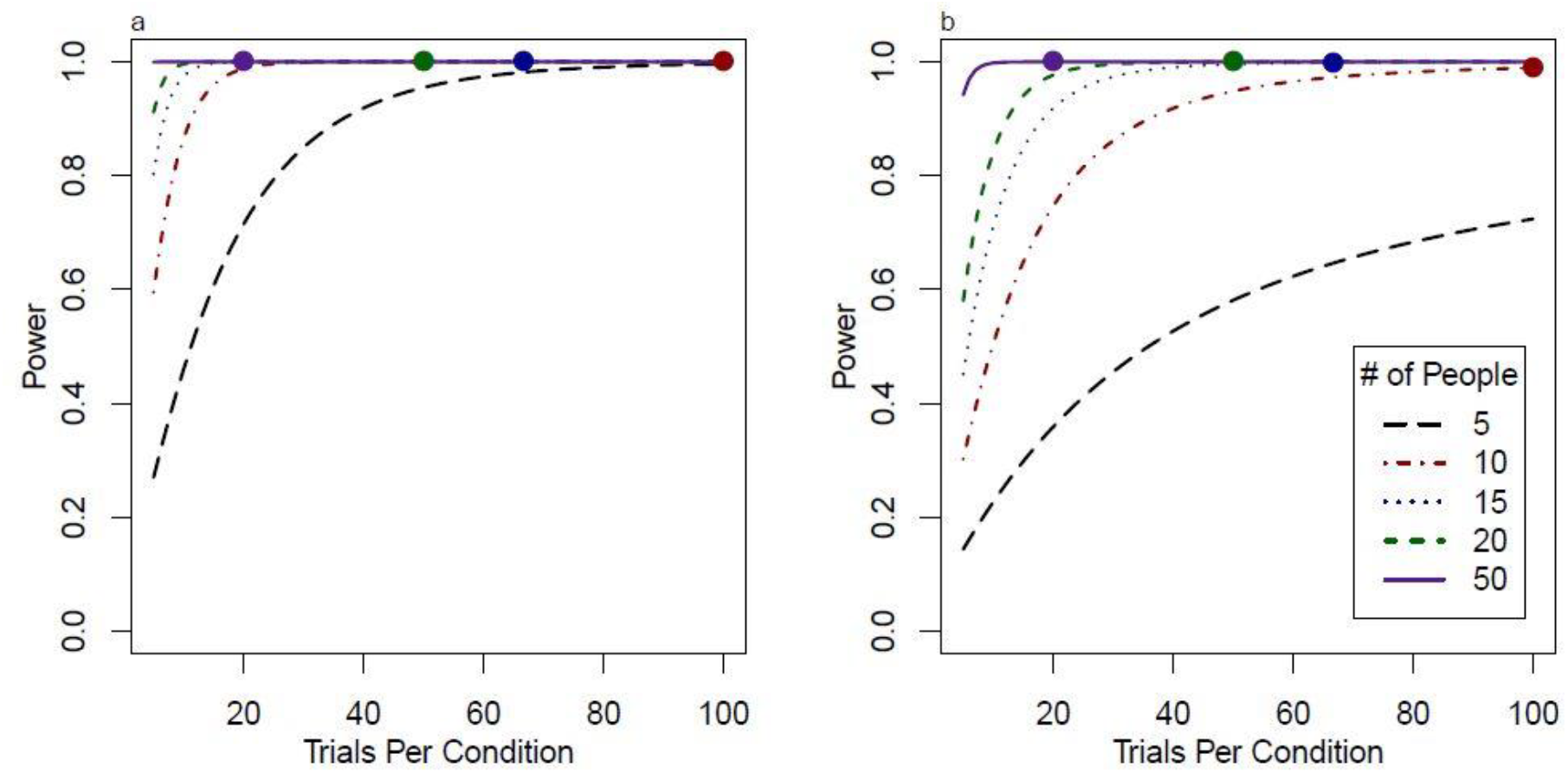
Power for different design scenarios. Panel 1A visualizes power for a very small effect (0.05 mean difference between gains) as a function of trials per condition for different numbers of participants. Panel 1B visualizes the same for a small effect (0.1 mean difference between gains). The calculations are based on the script https://github.com/PerceptionAndCognitionLab/stat-sampsize/blob/public/papers/amppsSub/fig-pow.R.

### Data Preparation and Analysis

#### Pre-registered Hypothesis I: Visual Gains

We used the Visual Gain, defined as v_target_/v_eyes_, as a proxy for quality of visual pursuit. To calculate the Gain from our data, we employed the following method, which you can also view in the R^35^ code we uploaded in our preregistration (https://osf.io/8vg95/; here, you can also find the final code that we used for data analysis to allow for a transparent comparison with the pre-registered code, as well as all data we collected in this experiment, and the additional code we used in the review process). With the surface tracker technology, the Pupil Labs eye tracking system saves the data in the real-world coordinates of the defined display. We first excluded all data points where the confidence reported by the pupil labs system was < 0.9 (6.4% of all frames), or where the gaze was outside of the display (y_eyes_ > 1 or < −0.3; x_eyes_ > 1 or < 0; 2.2% of the remaining frames). Furthermore, we excluded frames with missing data (0.1% of the remaining data). We then employed a series of computations to transform the data, which pupil labs reports in the coordinates of the display, into real world data of the visual scene, scaling the gaze positions on the surface to the depth of the stimulus. Therefore, gaze velocities are reported in terms of where the gaze hit in the z plane of the stimulus. We then smoothed the gaze position data locally with a Gaussian filter. Next, we calculate the tangential velocities with a lag of 9, and in a second step we calculate the acceleration as derivative of the velocity in time with a lag of 5. We defined as saccades those parts of the trajectory where acceleration was > 300 m/s^2^ and/or where the velocity of the eyes was > 1.5*velocity_target_. This is a relatively low value, but as Figure 3 shows, the algorithm identifies saccades with high accuracy. For each trial, we then calculated the average gain as v_tangential(eyes)_/v_tangential(target)_ for those parts of the trajectory (1) after the first catch-up saccade after motion onset, (2) that were not classified as saccades, (3) before the occlusion.

**Figure 3:**
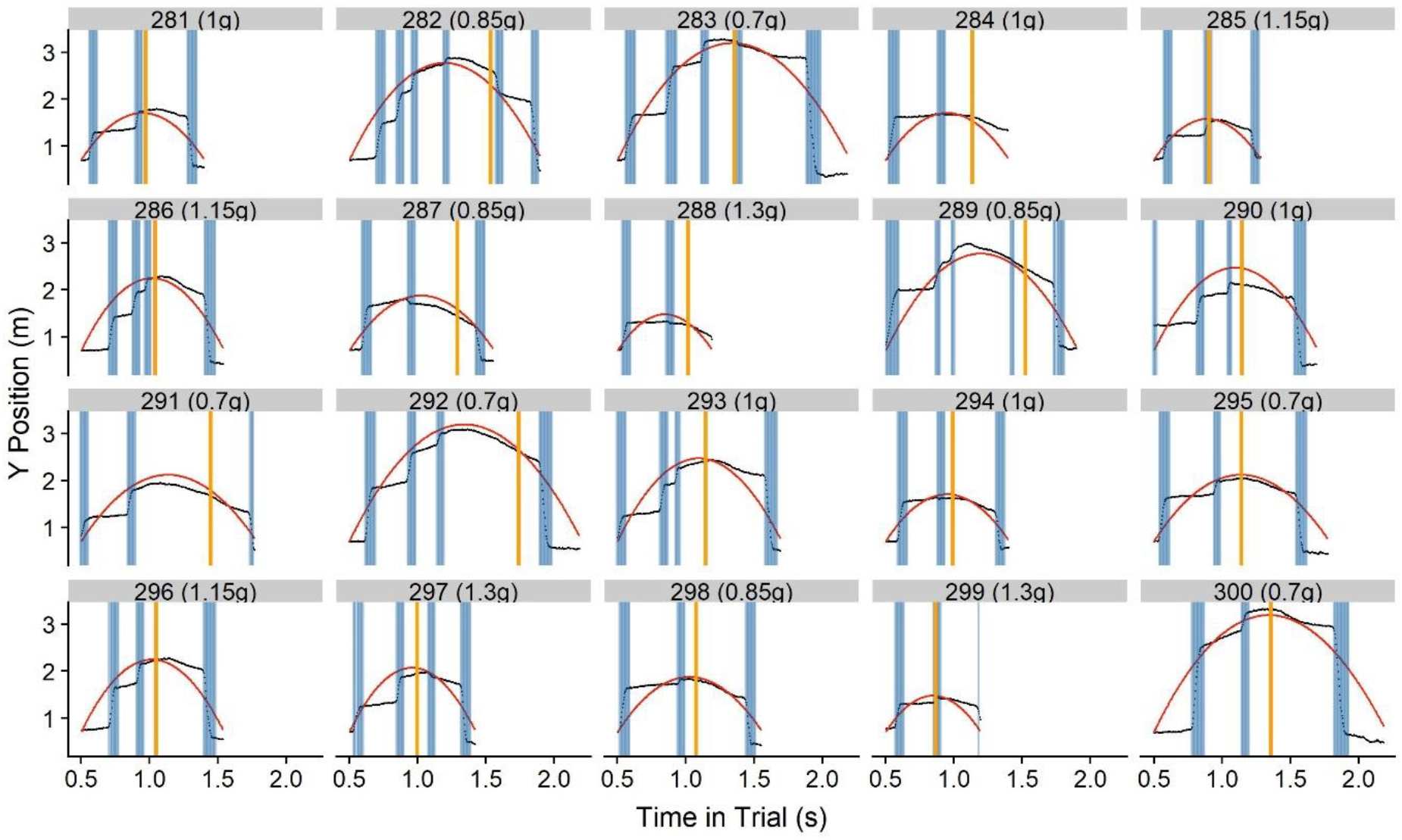
Eye behavior for 24 randomly chosen trials. The black dots indicate the recorded gaze behavior. The red line until the yellow vertical line is the actual trajectory of the ball. The red line after the yellow vertical line is the hypothetical trajectory the ball would follow after the occlusion. The vertical yellow lines indicate occlusion onset, while areas shaded in blue indicate behavior that our algorithm marked as saccades.

To test our hypothesis, we employed Linear Mixed Modeling on the pursuit gain as dependent variable to compare a model with subjects as grouping variable and gravity as fixed effect and intercepts as random effects to a Null Model. We analyzed the block where we presented 1g and −1g motion separately from the other blocks where we presented 0.7-1.3 g trajectories.

#### Pre-registered Hypothesis II: Timing Error

We exclude trials where the timing error was >= 0.5 s as outliers. Then, we employ Linear Mixed Modeling to compare a model with subject as grouping variable, Gravity, Occlusion Condition and their interaction as fixed effects and intercepts as random effects to a null model without Gravity as fixed effect.

In a further, non-preregistered effort, we establish a simple model to predict the timing error based on the last observed velocity in y direction, the missing distance and earth gravity. We also explore several post-hoc analyses that could grant insight into whether an observed effect may be due to a central tendency.

#### Exploratory Analysis: Saccades as indicator for spatial predictions

We furthermore explored the possibility that the first saccade after occlusion onset could serve as indicator of spatial predictions. To this end, we identified the x position of the gaze after the first saccade and subtracted it from the x position where the ball hit the table. A negative error thus means that the target location of the saccade was to the left of the point of coincidence, while a positive error means the saccade went to its right. We discarded the error in y direction because subjects had the table as a reference for the height, which should overrule any predictive information. We furthermore look at saccadic frequency, saccadic errors and time lag between gaze and target as potential further parameters of interest.

## Results

The hypotheses described under the subheadings “Visual Gains” and “Timing Error” were preregistered as confirmatory hypotheses, exceptions being the analysis where we compared bins of similar presentation times and an analysis that only included those subjects that had received the −1g/1g block first. Both were post-hoc analyses to identify or rule out potential confounds. The analyses described under “Modelling the Timing Error”, “Central Tendency?”, “Saccades as indicators of spatial predictions” and “Further Exploratory Analyses” are exploratory.

### Visual Gains

We compared a Linear Mixed Model (LMM) with subjects as grouping variable, gravity as fixed effect and intercepts per subject as random effects to a Null Model. For the 1g/-1g model, an ANOVA confirmed that the LMM was significantly better than the Null Model (p < 0.001). The difference in gain was −0.13 (SE = 0.005), confirming our hypothesis. Figure 4 visualizes the average gains per gravity condition for each subject. In a post-hoc, non-preregistered analysis, we only included data of subjects that received the −1g/1g block first, where both trajectories were presented with the same probability. Here, the difference in gain was even higher, with a mean difference in gains of 0.13 (SE = 0.005). This confirms that the effect was not based on 1g trajectories being presented more often throughout the experiment. Furthermore, the onset of smooth pursuit was on average only 3 m/s later than in the 1g trials, which provides further evidence that the disadvantages in pursuit gain for −1g trials was not merely due to the subjects’ surprise about the locus of appearance.

**Figure 4:**
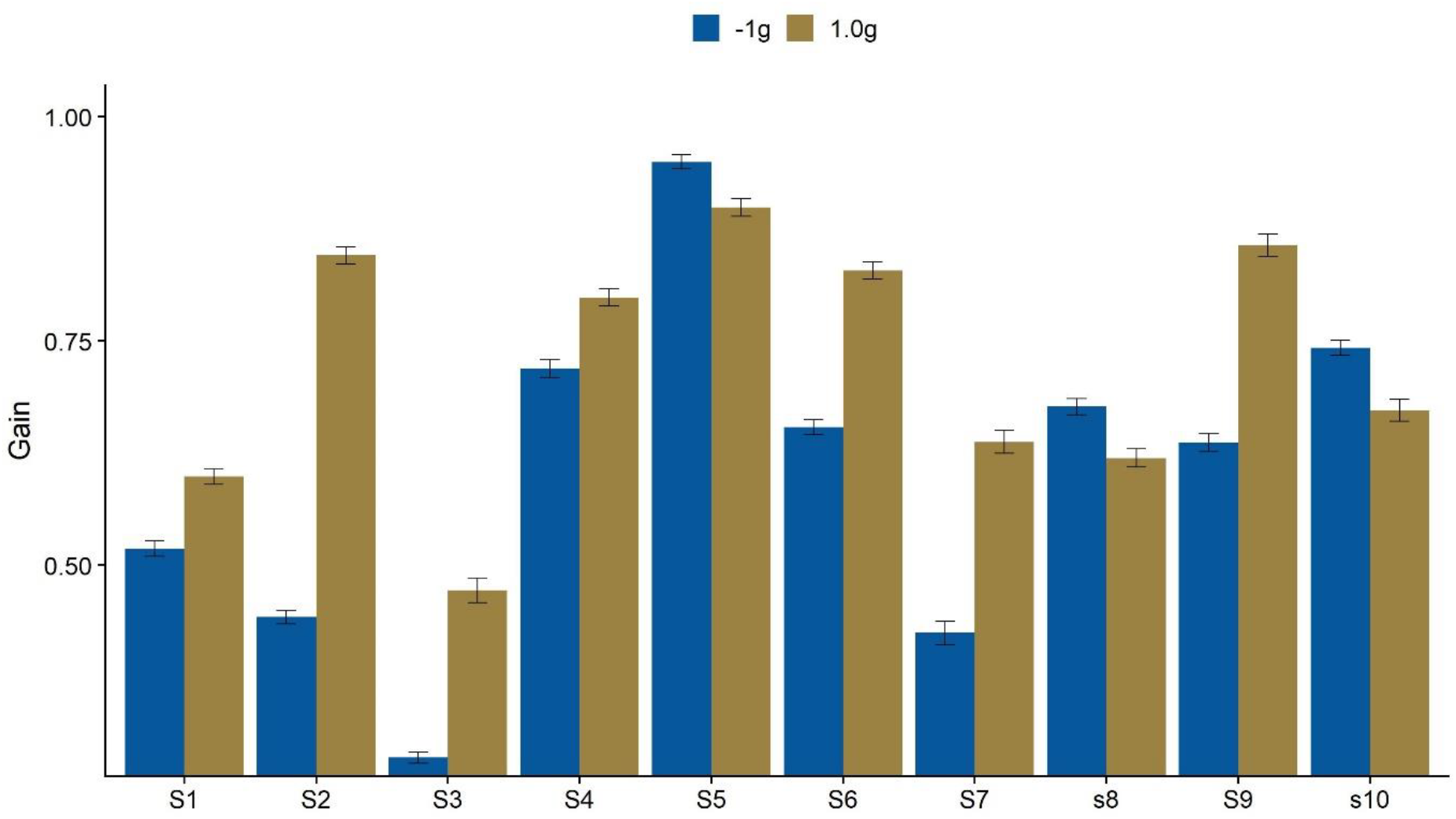
Average Gains for −1g and 1g per subject. Vertical lines on top of the bars indicate the standard error (+/− 1 SE). S1, S3, S5, S7 and S9 received the −1g block first. S2, S4, S6, S8 and S10 received the 0.7-1.3g block first.

For the 0.7g-1.3g model, an ANOVA confirmed that it was better than the Null Model (p < 0.001). However, the effect partially goes in the wrong direction: gain was higher for 0.7g (by 0.053, SE = 0.0054) and 0.85g (by 0.029, SE = 0.0054) than for 1g, and lower for 1.15g (by 0.043, SE = 0.0054) and 1.3g (by 0.065, SE = 0.0054), going against our hypothesis. The effect was much more homogeneous than for 1g/-1g; we thus visualize it grouped together for all subjects in Figure 5.

**Figure 5:**
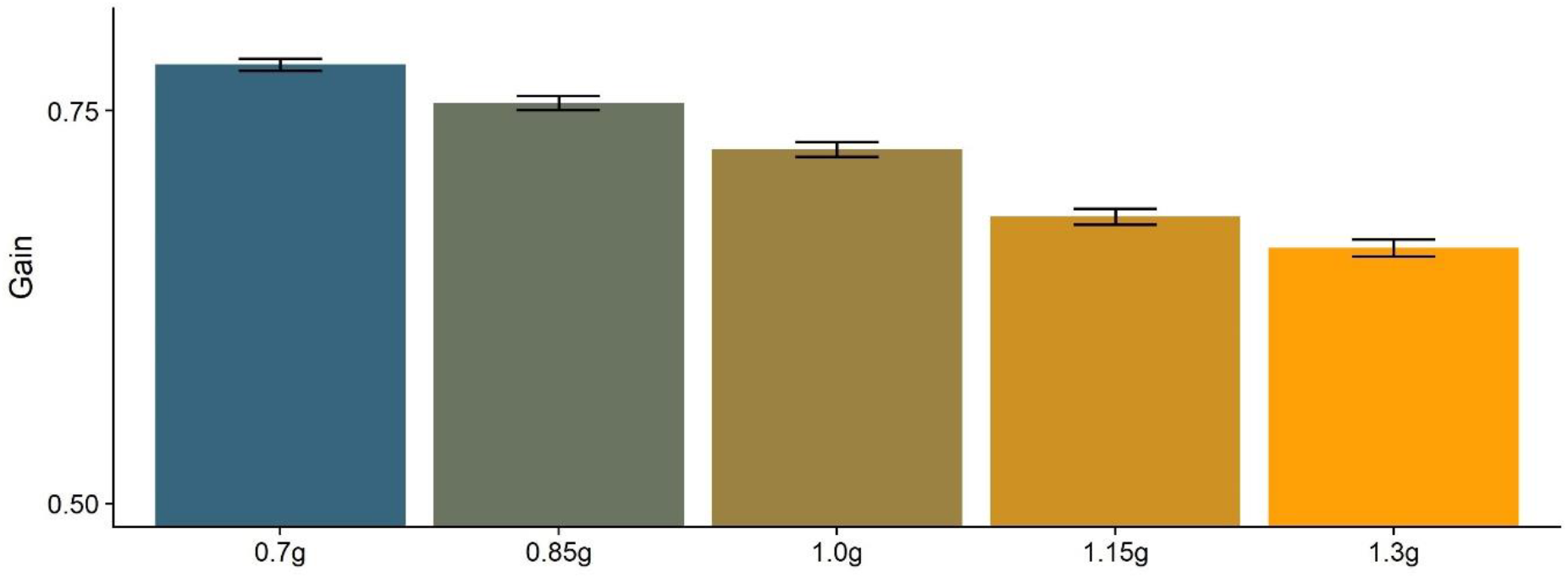
Gains averaged across subjects and gravity level. Lines on top of the bars indicate the Standard Error (+/− 1 SE).

Considering the unexpected result for the 0.7g-1.3g condition, we further analyzed whether this effect could be attributed to gravity, or if there were other factors at play. We discarded average speeds because they are virtually the same across gravities; mathematically, if the initial velocities are the same, the average velocity of trajectories with different gravities is the same for any given percentage range. We further tested whether differences in presentation time could lead to this effect. We used the variability we had in our presentation times due to the different occlusion times and initial velocities to compare trajectories with similar presentation times, but different gravities (see Figure 6). Within the bins, there still seem to be differences according to gravity levels; however, as different percentages of the trajectories are occluded, average velocities per trajectory necessarily differ, which may introduce a new bias.

**Figure 6:**
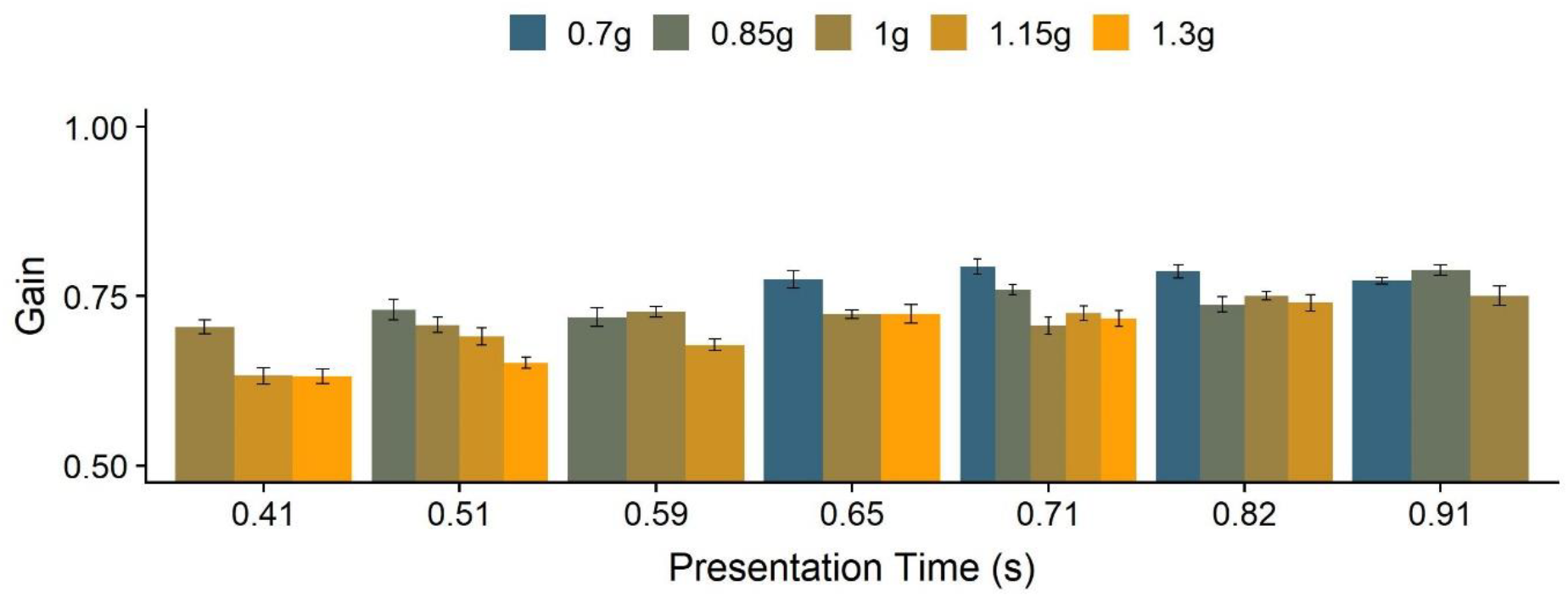
Gains for different presentation times, divided by gravity level. The variey in presentation times stems from the different combinations of gravity and initial vertical velocity, the Occlusion condition (long or short) and the variability in the occlusion length within each occlusion condition. We binned presentation times into seven bins with the same number of trials each, which resulted in the bins 0.44s, 0.51s, 0.59s, 0.65s, 0.72s, 0.73s and 0.82s. We calculate the average gains per gravity and presentation time as well as the Standard Errors; the black lines on top of the bars indicate +/− SE.

### Timing Error

We subtracted the mean system delay from the measured temporal errors. Furthermore, according to the pre-registration, we were going to compare a Linear Mixed Model (LMM) with subjects as grouping variable, Gravity, Occlusion Category and their interaction as fixed effects and intercepts per subject as random effects to a Null Model. However, for the 0.7g-1.3g condition, we found that a simpler model, with subjects as grouping variable, Gravity as fixed effect and intercepts per subject as random effects was not significantly worse than the model we had pre-registered (p = 0.11 for the 0.7g-1.3g model). For the −1g/1g condition, the original model was better than the simpler model (p < 0.001). We thus proceeded with the simpler model for the 0.7/0.85/1/1.15/1.3g condition and with the more complex model for the −1g/1g condition. For the 1g/1g model, an ANOVA confirmed that the LMM was significantly better than the Null Model (p < 0.001). For −1g, subjects pressed the mouse button later than for 1g, an effect that was stronger for the long occlusion condition (the regression line is given by Y = 0.04*Gravity Category+0.005*Occlusion Category+0.02*Gravity Category*Occlusion Category+0.1), confirming our hypothesis. Figure 7 visualizes the average timing errors across all participants per gravity condition.

**Figure 7:**
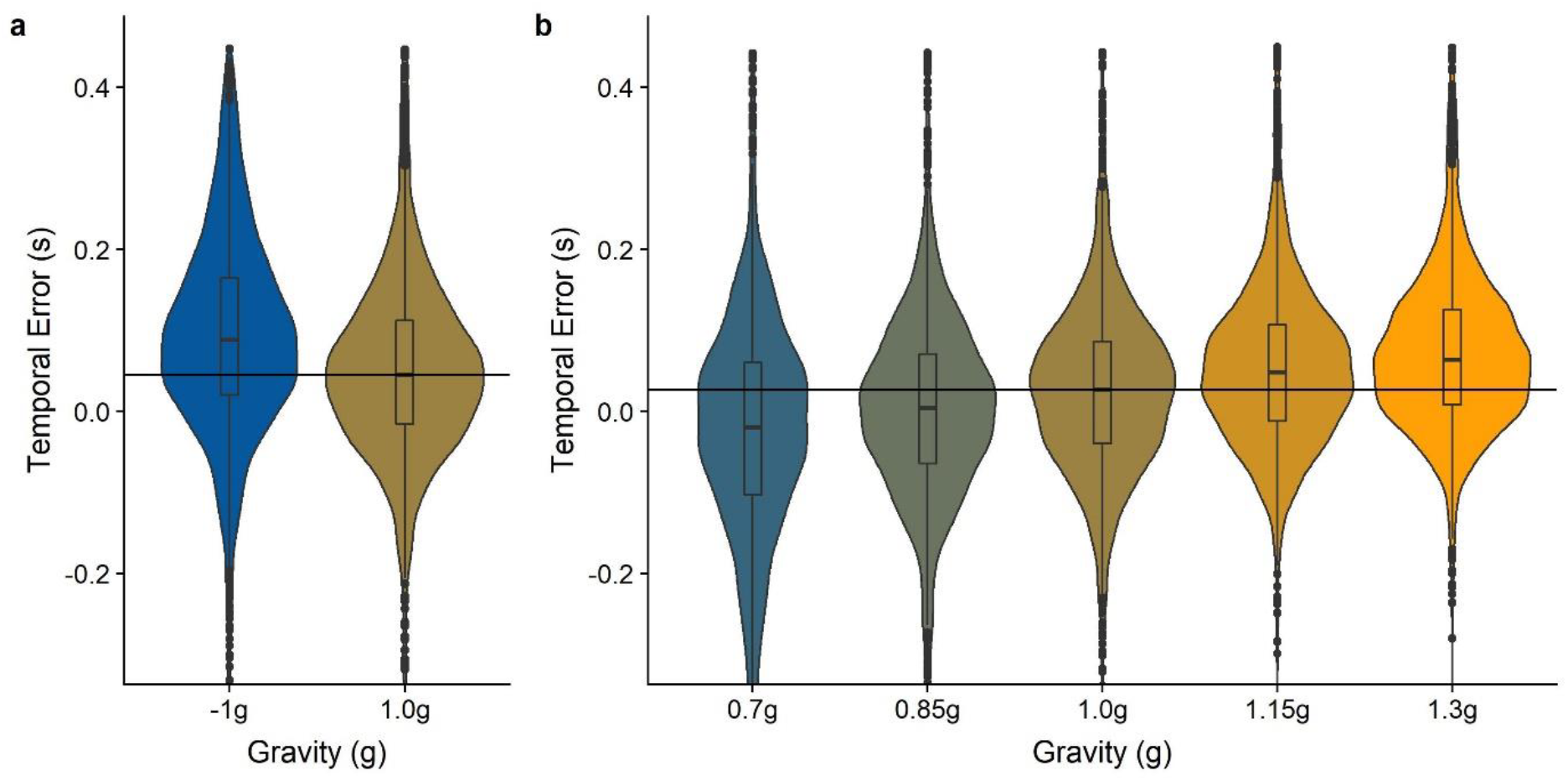
Temporal errors per gravity condition. Negative values indicate that subjects pressed the button too early with regards to the moment of impact. Positive values indicate that the button was pressed too late. Boxplots in the middle of each structure indicate the percentiles and the median, while the wings represent the distribution of responses. The horizontal black line represents the median of the responses for 1g. Figure 6a visualizes data from the −1g/1g blocks, while Figure 6b visualizes data from the 0.7/0.85/1/1.15/1.3g blocks. This figure shows temporal errors after subtracting the mean system delay.

For the 0.7g-1.3g model, an ANOVA confirmed that it was significantly better than the Null Model (p < 0.001). Subjects pressed the mouse button earlier for 0.7g (by 0.05 s, SE = 0.0035 s) and 0.85g (by 0.024 s, SE = 0.0035 s), and later for 1.15g (by 0.023 s, SE = 0.0035) and 1.3g (by 0.040 s, SE = 0.0034 s), confirming our hypothesis. See Figure 7 for a visualization of the differences.

### Modelling the Timing Error

For further analysis of the temporal error, we excluded subject s09 because they displayed a mean temporal error way above the mean of the remaining subjects (mean error = 0.23 s, while the other subjects displayed mean temporal errors of −0.08 to 0.1 s). Both the mean temporal error for 1g in the 0.7/0.85/1/1.15/1.3g condition (mean = 0.018 s, p < 0.001, t = 7.6639) and in the −1g/1g condition (mean = 0.04 s p < 0.001, t = 16.287) are significantly different from zero, even after excluding subject s09 and accounting for the delays measured in the projectors. The effect was stronger for the short occlusions (mean temporal error for long occlusions = 0.014 s, mean temporal error for short occlusions = 0.046 s, p < 0.001, t = −15.125).

To verify to what extend the observed temporal errors are consistent with the use of an earth gravity prior to extrapolate motion during the occlusions, we simulated the temporal errors assuming the last vertical velocities observed by the participants, a gravity value of 9.81 m/s^2^ and the remaining distance in y direction. This model is based on the physical formula for travelled distance from initial velocity and acceleration (Equation 3), solved for time (Equation 4).

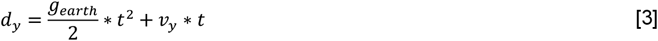

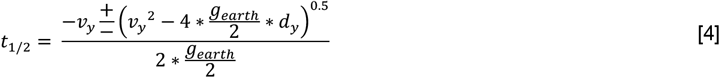

d_y_ is the remaining distance in y direction, g_earth_ is the value earth gravity (9.81 m/s^2^) and v_y_ is the last observed velocity in y direction. As we are interested in future positions, we discard negative solutions of Equation 4 for t (that is, the version of the formula where the discriminant is subtracted). We did not consider air drag here, as the timing differences are minuscule (e. g. 1.5 ms for 1g, long occlusion and 6 m/s initial vertical velocity), while adding air drag to the equations would lead to an exorbitant increase in mathematical complexity (see Equations 1 and 2).

Figure 8a depicts the observed errors as a function of the error predicted from our simple model. The overall correlation between simulated and observed temporal error was r = 0.31, while it rose to r = 0.37 when only considering the long occlusion condition and dropped to r = 0.17 for the short occlusion condition. This indicates that gravity becomes more important as a source of information as the occlusion becomes longer and motion is predicted for a larger time frame. In Figure 8B, we illustrate the median simulated error and the median observed error for the two occlusion conditions and each gravity value. As apparent from the plot, the observed data fit the predictions very well for the Long Occlusion condition, indicating that humans indeed rely on earth gravity to guide temporal coincidence responses. The predictions for the Short Occlusion condition fit the trend of the observed errors well, while participants’ responses are consistently too late. This systematic temporal error cannot be explained to delays in our system, and should represent a veridical reaction of the participants to our stimulus. A possible explanation is that the time of coincidence with the table (mean of 0.3 s across all participants and conditions after occlusion) coincides roughly with the time of the first saccade after occlusion (mean of 0.21 s across all participants and conditions after occlusion). Participants may either need some time to monitor information after the saccade, or the planning of the motor command to execute this saccade interferes with the planning of the motor command to initiate the click.

**Figure 8:**
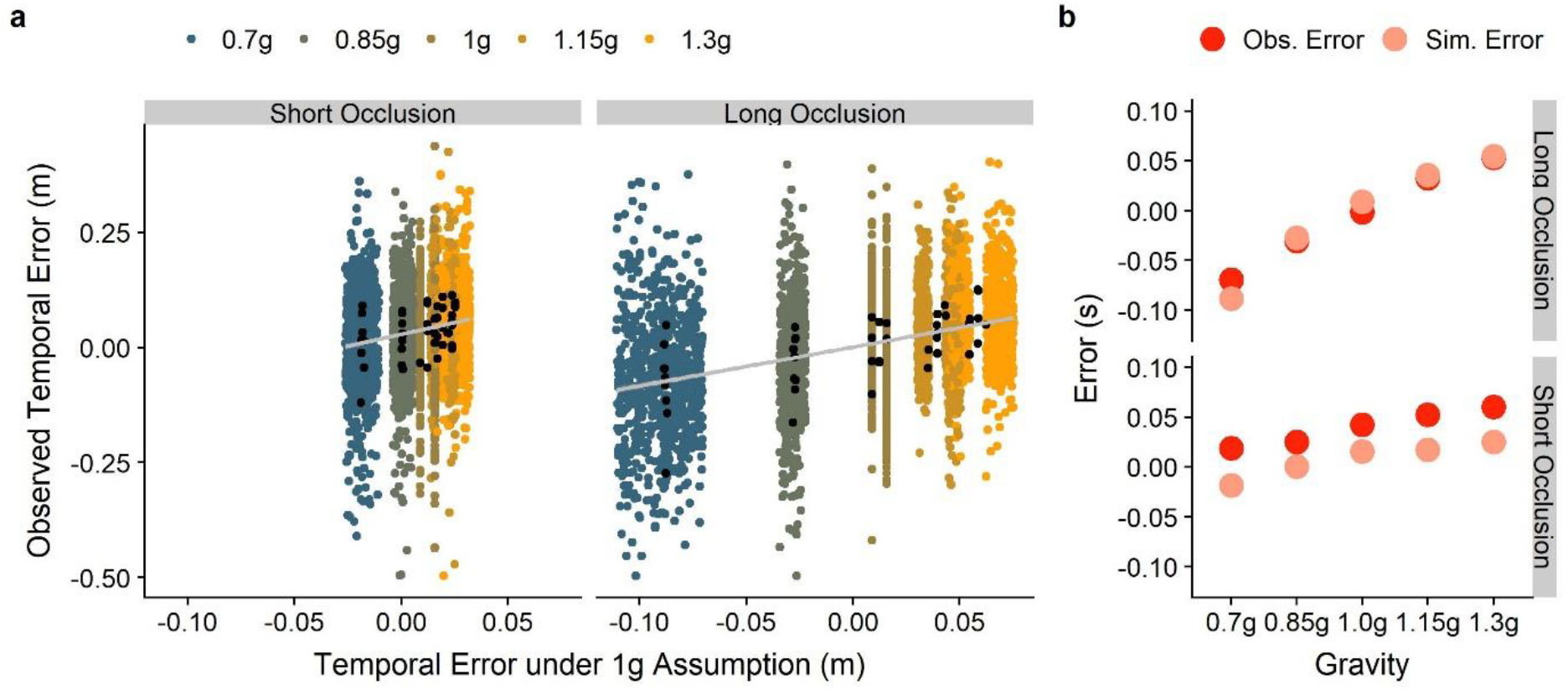
A. Empirically observed temporal errors as a function of the error predicted by our model. We display the data separately for the short and the long occlusion conditions and color coded for gravity. The black dots are the median values per subject. The grey line is the regression line, while the (very) faint grey area around it marks +/− 1 standard error. Due to the amount of data points, the standard error is extremely small and barely visible in the plot. B. Median observed errors per gravity condition (red) and median simulated errors per gravity conditions (light red).

### Central Tendency?

As the results for 0.7-1.3g are roughly symmetric, it could be argued that our timing results are due to a regression to the mean. Two reasons speak against this interpretation: First of all, the dispersion differs across conditions. If our results were due to a central tendency, we would expect roughly the same variability across all TTCs. However, dispersion increases with decreasing gravities, most likely due to higher TTCs, as depicted in Figure 9a. Secondly, the variability in our stimulus (two different initial vertical velocities) allows us to compare different TTCs within one gravity level. If there was a central bias in subjects’ responses, then both should be biased heavily towards their common mean. In Figure 9b, we show the mean extrapolated times as indicated by subjects’ responses (“Response”) and the actual duration of occlusion (“Stimulus”). We observe that the difference in responses is slightly smaller than the actual difference in the stimuli. Thus, there *may* be a regression to the mean, however, its effects are tiny in comparison to other processes at play. As a third piece of evidence, a central tendency and a 1g assumption lead to numerically different predictions about the distribution of errors. In Figure 9c, we disentangle these predictions: While the general bias to answer too early in the Short Occlusion condition makes it hard to draw conclusions whether a central bias or a 1g assumption is responsible for the observed patterns, the picture is much clearer in the Long Occlusion condition: if subjects used a mean value of all response times to time their responses (black dots), we would expect a systematic bias to answer too early. The observed behavior (solid red dots) match much more closely the predictions of our 1g model (translucent red dots).

**Figure 9.**
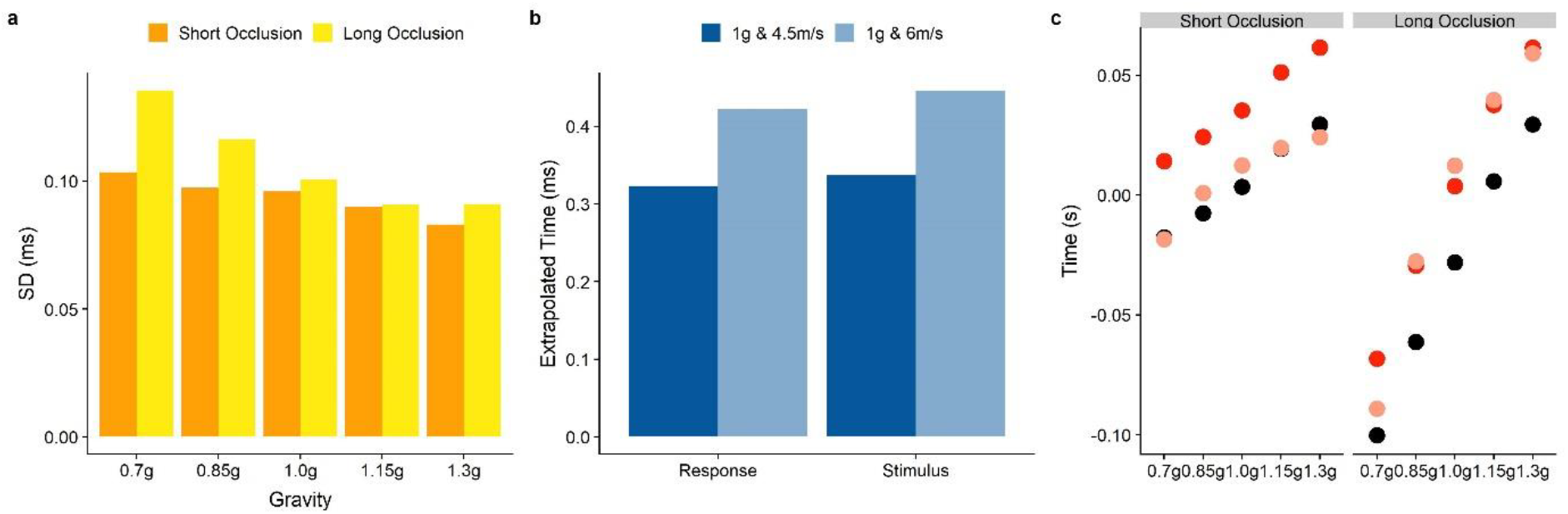
A. A comparison of standard deviations across the different gravity conditions, separated by Occlusion Condition (orange for short and yellow for long). B. Response: Mean extrapolated motion duration during occlusion, as indicated by the participants responses, with extrapolated time being defined as occluded time + timing error. Stimulus: Duration of occlusion. Dark blue are the values for an initial vertical velocity of 4.5 m/s and a gravity of 1g, while light blue indicates trajectories with 6 m/s and 1g. C. The predictions of the 1g assumption model (light red), predicted responses under a central tendency (black dots) and actual responses (red).

### Saccades as indicators for spatial predictions

Apart from the two pre-registered hypotheses, we tested whether the first saccade after occlusion onset would reveal gravity-based spatial predictions. This error was computed as target x position of the first saccade after occlusion onset minus the position where the ball hit the table (see also code available under https://osf.io/8vg95/). We compared a Linear Mixed Model (LMM) with subjects as grouping variable, Gravity as fixed effects and intercepts per subject as random effects to a Null Model. As we did not expect any spatial bias for −1g, we applied this model only to the 0.7-1.3g condition. An ANOVA showed that the model was significantly better than the Null Model (p<0.001). In comparison to 1g trajectories, subjects executed their first saccade after occlusion onset more to the left for 0.7g (0.77 m, SE = 0.042 m) and 0.85g (0.29 m, SE = 0.042 m) trajectories, and more to the right for 1.15g (0.12 m, SE = 0.042 m) and 1.3g (0.29 m, SE = 0.042 m) trajectories. This effect is consistent with assuming 1g for the occluded part of the trajectory. Figure 10 visualizes this effect.

**Figure 10:**
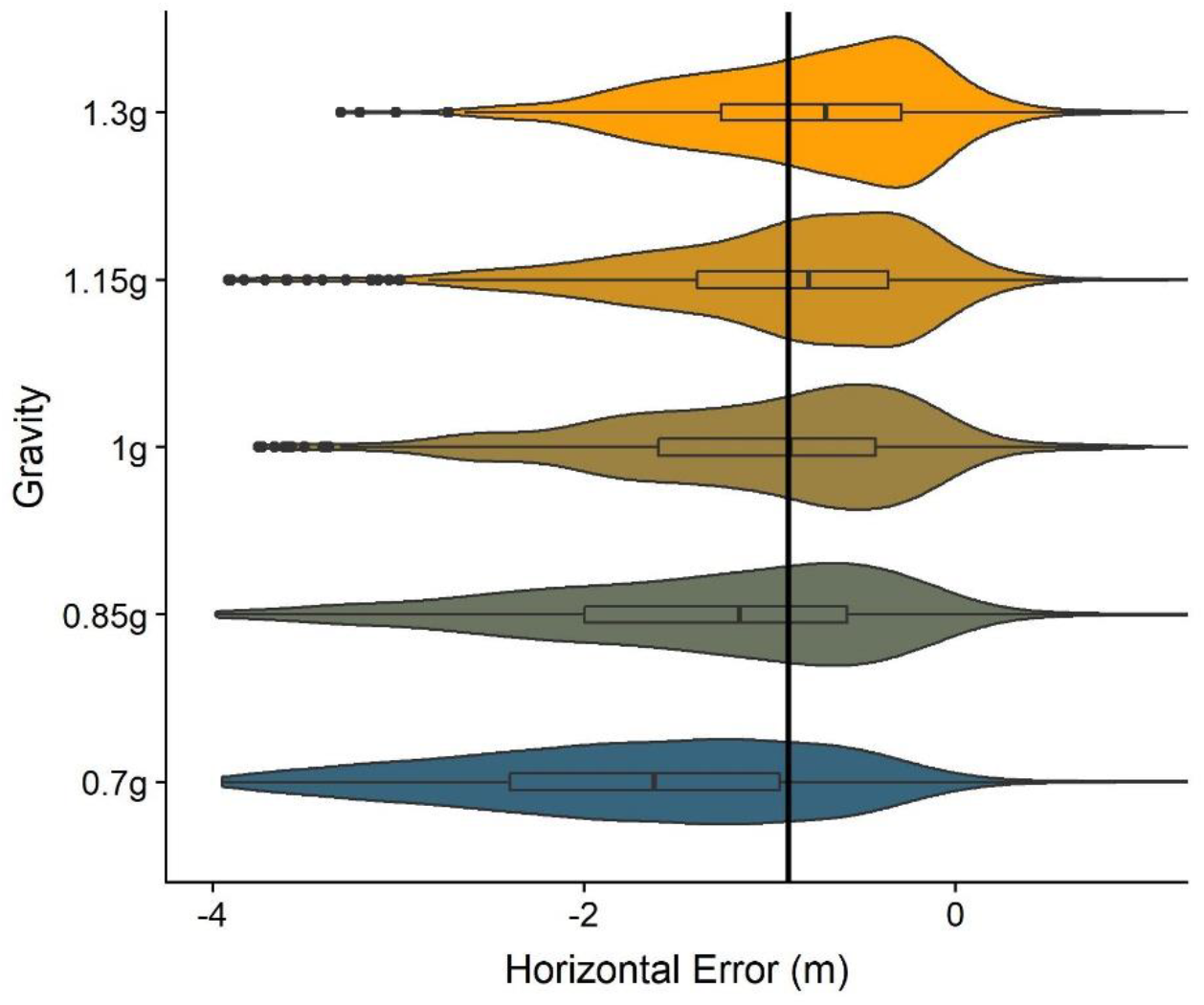
Spatial Error per gravity level. Negative values represent an undershoot, that is, the saccade lands further to the left than the spot where the ball hits the table. Positive values in turn signify an overshoot. Boxplots in the middle of each structure indicate the percentiles and the median, while the wings represent the distribution of responses. The black line represents the median of responses for 1g.

### Further Exploratory Analyses

Last but not least, we conducted further exploratory analyses on parameters that might provide additional evidence. Among these parameters are the lag between eyes and target, the number of saccades and saccadic errors. In Table 1a you find the means and standard errors of the respective values for the −1g/1g condition. (1) For the lag between eyes and target, we compared results from −1g and 1g trials. We calculated the spatial error between eyes and target at 40 % of the trajectory as an indicator of the overall lag between eyes and target. Unintuitively, we found that the x error was bigger for 1g than for −1g. Upon further analysis, it becomes clear, however, that predictions in x direction are independent of gravitational information, and may be established based on the observed velocity in x direction alone. On the other hand, in line with our general hypothesis that gravitational motion should facilitate eye-movements, the y error was smaller for 1g than for −1g. (2) We found that 1g trials elicited more saccades than −1g trajectories. This is a slightly caveat for our overall conclusions, as less predictable motion should lead to a higher saccadic frequency. (3) We used the error after the first catch-up saccade as surrogate for the overall saccadic errors and found that the error in both x and y direction was smaller for 1g than for −1g.

**Table 1a:** Values and standard errors for different parameters that could be used to assess the use of prediction information for eye-movements. Negative values mean that the gaze was directed to the right of the target, or below the target, respectively. This table covers the-1g/1g condition.

| Parameter | 1g | -1g |
| --- | --- | --- |
| Lag between eyes and target: x | -0.34±0.01 m | -0.25±0.02 m |
| Lag between eyes and target: y | -0.40±0.01 m | -0.45±0.02 m |
| Number of saccades | 2.28±0.04 | 1.94±0.03 |
| Saccadic error: x | -0.45±0.01 m | -0.53±0.01 m |
| Saccadic error: y | -0.63±0.01 | -0.73±0.01 m |

Table 1b shows the results for the 0.7g/1.3g condition. For the lag, we chose 0.4 s after motion onset as point of reference instead of 40 % of the trajectory, as 40 % of the trajectory may coincide with, or by influenced by, the first catch-up saccade for trajectories with the highest gravities. 0.4 s into the trajectory, the first catch-up saccade was finished in most trials and smooth pursuit had already been initiated. For this parameter, we find that lag in both x and y direction decreases with increasing gravity. For the y direction, this is in line with our general hypothesis that eye-movements are partially guided by an internal model of earth gravity: subject expect the target to gain height faster than it actually does for higher gravities, making their gaze trail the target less than for 1g motion, and vice-versa. The result is less intuitive for the lag in x direction, as predictions in the x direction should be based only on the horizontal velocity. This may indicate that an internal representation of earth gravity may be used even to recover the physical horizontal velocity from optic flow input (as would be computationally useful^21^). With regards to the saccadic errors, we find a similar, but less pronounced pattern, to which the same rationale applies as to the lag between gaze and target.

**Table 1b:** As Table 1a, but for the 0.7g-1.3g condition. Note that we did not include the number of saccades for this condition, as the differences in motion duration make a comparison virtually meaningless.

| Parameter | 0.7g | 0.85g | 1g | 1.15g | 1.3g |
| --- | --- | --- | --- | --- | --- |
| Lag: x | -0.50±0.02m | -0.43±0.01m | -0.37±0.01m | -0.35±0.01m | -0.33±0.01m |
| Lag: y | -0.50±0.01m | -0.46±0.01m | -0.41±0.01m | -0.35±0.01m | -0.26±0.01m |
| Saccadic error: x | -0.48±0.01m | -0.44±0.01m | -0.41±0.01m | -0.41±0.01m | -0.41±0.01m |
| Saccadic error: y | -0.57±0.01m | -0.55±0.01m | -0.54±0.01m | -0.52±0.01m | -0.51±0.01m |

Last but not least, we checked whether ocular pursuit performance would predict the timing error. It stands to reason that a closer pursuit between peak and occlusion could enhance the representation of the last observed velocity before disappearance. A representation of this velocity is one of the terms in our 1g model for the temporal error, and a more precise representation of this value could lead to more precise predictions. However, the velocity term plays a subordinate role with regards to the gravity term; we thus expect any influence to be relatively modest. Its weighting is somewhat higher for late occlusion trials, where a slightly higher impact is expected. To address this question, we correlate the average gain for the time window between peak and disappearance with the absolute value of the timing error in the same trial, across all blocks and participants, but separately for early and late occlusion trials. For early occlusion trials, we find no correlation to speak of (r = 0.006, p = 0.642, t = 0.46). For late occlusion trials, we found a tiny, non-significant correlation (r = 0.02, p = 0.096, t = 1.66).

## Discussion

Our results show that humans use an earth gravity prior to guide their smooth pursuit eye movements, and possibly also to guide saccades. This effect is clearly visible in the 1g/-1g condition, which we consider to be the stronger test of our hypothesis as it minimizes confounds. The results from the 0.7g-1.3g condition are less clear: we observed a correlation between gravity value and gains, which goes against our original prediction that the highest gains should be observed for 1g trajectories, with lower gains for both lower and higher gravities.

An alternative is a post hoc adjustment of the original hypothesis – which, for this reason, should be treated with the due caution: Since the gains are calculated to the bigger part of ascending trajectories, employing earth gravity to guide smooth pursuit would mean that the gaze is lagging behind for higher gravities, and slightly ahead of the target for lower gravities. This is exactly the direction of effect we found in the data. However, if taken seriously, this interpretation would mean that gains should be around 1 for 1g, below 1 for 1.15 and 1.3g and above 1 for 0.7 and 0.85g. As gains are generally below 1, our data can’t be taken as strong evidence for this interpretation of the original hypothesis. Also, after further analyzing the data, we believe that the effect is an artifact of our design, where higher gravities entailed lower presentation times. Lower presentation times could impact pursuit because subjects have less time to adjust to the trajectory. It is, unfortunately, virtually impossible to match every relevant dimension of the stimulus: to match the presentation time while maintaining gravity, for example, the initial vertical velocity would have to be adjusted, which may incur new problems.

Our results also shed light on what information the visual system relies upon to make predictions about observed motion in order to guide eye-movements. Generally, it has been found that motion estimates are taken into account to predictively guide eye movements^36,37^: wave forms are pursued more closely than neural delays would allow without prediction^38^ and curvilinear paths were followed throughout an occlusion^39^. Arbitrary accelerations are taken into account qualitatively at best^18,19,40^. The control of eye movements has also been described in an active inference framework^41,42^, and neural bases have been suggested^43,44^ for predictive mechanisms. We identified three studies that are directly relevant to the theme of the present paper: Souto & Kerzel^45^ showed that congruency of translational and rotational motion of an object aided ocular pursuit, taking advantage of the fact that only the congruent motion was possible in the real world. However, the relationship with gravity is quite intermediary in this case, as it only served to constrain which of the scenarios were congruent with reality. It is therefore not clear whether this effect can be attributed directly to gravity, or more generally to a real-world feasibility of the stimulus. Furthermore, Delle Monache et al.^22^ investigated eye-movements in response to 0g and 2g perturbations in parabolic 1g motion in front of a pictorial background. They found that tracking was enhanced for 2g with regards to 0g, showing that a qualitative expectation of downwards moving objects to accelerate is also present for eye behavior. A follow-up study from the same group^26^ showed similar results when motion was presented in front of a pictorial background, but no clear signs of prediction dependent ocular behavior when motion was presented in front of a uniformly black background. Also Kreyenmeier et al.^28^ highlight the need to take the context of motion into account, at least for ocular pursuit in response to visual stimuli. These results highlight the need for realistic stimuli for the study of gravity-based prediction in eye-movements. While our results are generally in line with both studies by Delle Monache et al., our stimuli contain not only pictorial cues, but also 3D information, which makes them the most ecologically valid in this line of research. By demonstrating that congruency with earth gravity improves visual pursuit gains we show that, in addition to the aforementioned sources of information, also an earth gravity prior is recruited to guide eye movements.

Finally, differences in gains for 1g and −1g motion could be due to anisotropies with regards to motion direction. However, Ke et al^46^ found that gains were slightly lower in response to diagonal left-down to right-up motion – which, due to the parts of the trajectory we counted towards gains, corresponds roughly to our 1g stimulus – than in response to diagonal left-up to right-down motion (−1g). We would thus expect *higher* gains for −1g if this anisotropy was the only effect at play.

We furthermore aimed to replicate the known effect that internalized knowledge of gravity is used to extrapolate motion information for interception tasks (pre-registered hypothesis II). Our data support this notion: subjects generally clicked too late for higher gravities and too early for lower gravities (in the 0.7g-1.3g blocks), and too late for −1g, indicating that they expect objects to accelerate/decelerate according to earth gravity.

Moreover, to our knowledge, this is the first time a continuous, quantitative effect of an internal model of gravity has been shown. Studies have generally focused on comparing 1g to 0g or −1g motion to tease out the impact of our knowledge of gravity^8,25,47,48^. By analyzing the effect across a range of values (0.7/0.85/1/1.15/1.3g), we add an understanding of how our internalized representation of 1g affects coincidence timing across a range of gravity values. To help capture this quantitative aspect, we present a very simple model that estimates the temporal error based on the last observed vertical velocity before occlusion, the remaining distance and earth gravity. The model explains responses well, which can be taken as additional evidence that humans maintain and access a representation of gravity that is very accurate and nearly immutable. It furthermore stresses the quantitative nature of these gravity-based predictions, in comparison to previous results^2,3,6–8,22,26,49–51^, which have only shown qualitative effects.

The symmetry of the timing results suggests that there may be a central tendency or regression to the mean at play. However, when delving deeper into the data, there are several reasons that speak against this interpretation: First of all, the variabilities differ among the gravity levels, while a regression to the mean should lead to a uniform dispersion. Second, when comparing different TTCs within one gravity level, we find that the difference in estimation is only slightly smaller than the difference present in the stimulus. A regression to the mean would lead to no or much smaller differences. Finally, a regression to the mean and our earth-gravity based model yield numerically different predictions; overall, our data fit the 1g model better than a central tendency model.

When analyzing the data, we realized that many subjects made a saccade right after the target disappeared. As exploratory analysis, we verified whether also this saccade may be informed by earth gravity. Figure 10 illustrates that this may well be the case, and our LMM analysis confirmed that the data support this interpretation. In our data set, the effect is much stronger for lower gravities than for higher gravities, which might reflect a confound: for our stimuli, lower gravities meant that the point of coincidence was much further to the left than for higher gravities. Therefore, also the distance that needed to be covered during the saccade was much higher, resulting in an additional undershoot for higher gravities. In fact, a similar behavior has been shown for smooth pursuit^39,52^, where during a transient occlusion the gaze decelerated and stabilized at a lower velocity, lagging behind the hypothetical path of the target. In occlusion scenarios, saccades seem to be executed to compensate for lagging smooth pursuit, realigning the gaze with the position of the target^53^. Also independently of smooth pursuit, humans directed saccades to the expected position of a target after an occlusion^54^. We thus believe that the overall undershoot we are observing for all conditions is not a general trend of saccadic behavior for occlusions, but rather a consequence of our experimental design. All in all, especially because lower gravities required bigger saccadic amplitudes and thus more energy to reach the locus of coincidence, the data should be treated as suggestive rather than conclusive. Nonetheless, a further exploratory analysis on saccadic errors lends further support to this tentative hypothesis: the error after the first catch-up saccade after motion onset (in the ascending part of the trajectory) was smaller for 1g than for −1g. Similarly, in the 0.7g-1.3g condition, we found that errors were smaller for higher gravities and vice-versa. This could be due to subjects expecting the target to gain more elevation than it actually does for higher gravities, and less elevation for lower gravities. The notion that not only smooth pursuit, but also saccadic eye-movements are partially guided by an earth gravity prior, deserves thus further, confirmatory research.

Finally, we found no correlation between tracking performance before occlusion and performance in the timing task. This is partially in dissonance with results reported by Fooken et al.^27^: in a task structured similarly to ours, participants were asked to pursue a target on a linear trajectory. An occlusion ensued, after which participants had to intercept the target with their finger in a hitting zone. There may be different reasons for the discrepancy between this finding and our result: First of all, Fooken et al. employed linear motion in a 2D environment, while we employed parabolic motion in a 3D environment. The resources used to predict motion may differ considerably between these two contexts; several studies on the role of an internal model of gravity, for example, have shown that the 1g model is only recruited for naturalistic stimuli (e. g. pictorial background, 3D presentation)^23,26,49^. Online information might thus be more important in less naturalistic tasks, while more naturalistic tasks recruit models more strongly that are fed by prior knowledge about the real world. Furthermore, the final velocity of our target, which is arguably the kinematic component whose perception stands most to benefit from a more accurate pursuit, had a minor contribution to the final timing error in comparison to the effect of gravity. In line with this reasoning, the correlation between tracking and timing performance we observed was slightly higher (although still negligible) where the velocity component had a higher impact on the final timing. Finally, our participants had much more time to track the target and gather information about its kinematics than Fooken et al.’s participants. A high pursuit accuracy may therefore not have been necessary because the long duration of visible motion gave them enough time to establish a precise-enough representation of the relevant visual parameters. More in line with our results, Cesqui et al.^24^ found that the only ocular pursuit parameter to reliably predict catching performance for targets approaching frontally on a parabolic trajectory was the overall tracking duration – a parameter that we can’t assess in our design as subjects were instructed to follow the target with their gaze from beginning to disappearance. Just like us, Cesqui et al. found no clear population-wide correlation between catching performance and any other ocular parameters. This study differed from ours in that the stimulus moved in the sagittal plane and targets were caught manually, while our stimulus moved in the fronto-parallel plane and the coincidence was timed per button press. All in all, both Cesqui et al.’s and our studies suggest that there is no clear, generalizable relationship between gaze behavior and interception/coincidence timing for naturalistic 3D motion.

## Conclusions

In our study, we test the hypothesis that an earth gravity prior is used by humans to guide their smooth pursuit eye movements. We furthermore replicated a known effect where this earth gravity prior lead to systematic timing errors for interceptive timing. This effect had so far been studied only qualitatively, on which we add by demonstrating that a larger deviation from earth gravity leads to larger timing errors. Moreover, in exploratory analyses, we show that also saccades may be guided by earth gravity-based predictions.

Future research should confirm that saccades are guided by the earth gravity prior. More ambitiously, it would be interesting to delve deeper into quantitative aspects of the internal model of gravity: Do the introduced biases get stronger the further we are moving away from 1g? Are subjects able to adjust manual or ocular responses when given feedback and enough opportunity to learn? Is it, generally, more useful to envision our representation of earth gravity qualitatively as an internal model, or should we call it a gravity “prior”, invoking a Bayesian framework of perception and action?

## Author Contributions and Notes

BJ conducted and analyzed the experiment and wrote the paper. JLM provided the initial research question and provided overall advice. BJ and JLM programmed the stimuli in a joint effort.

The authors declare no conflict of interest.

## Acknowledgments

Funding was provided by the Catalan government (2017SGR-48) and the project ref. PSI2017-83493-R from AEI/Feder, UE. The first author (BJ) was supported by an FI fellowship (FI-DGR 2016) from the Catalan government.

## Notes

#### Summary of Updates

We have updated the article in response to very helpful commentary from our two reviewers in the revision process at Scientific Reports.

https://osf.io/8vg95/

